# Cerebral Autoregulation, CSF outflow resistance and outcome following CSF diversion in Normal Pressure Hydrocephalus

**DOI:** 10.1101/223867

**Authors:** Afroditi Despina Lalou, Marek Czosnyka, Joseph Donnelly, John D. Pickard, Eva Nabbanja, Nicole Keong., Matthew Garnett, Zofia Czosnyka

**Author notes:** Afroditi Despina Lalou, Newnham College, University of Cambridge, CB3 9DF. Disclosure: The patient data are derived from our ICM+ data archive and could possibly partially overlap with those of previous studies on NPH.

## Abstract

**Introduction:** Normal pressure hydrocephalus (NPH) is not simply the result of a disturbance in cerebrospinal fluid (CSF) circulation, but often includes cardiovascular comorbidity and abnormalities within the cerebral mantle. In this study, we have examined the relationship between the global autoregulation pressure reactivity index (PRx), the profile of disturbed CSF circulation and pressure-volume compensation, and their possible effects on outcome after surgery.

**Materials and methods:** We studied a cohort of 131 patients, investigated for possible NPH. Parameters describing CSF compensation and circulation were calculated during the cerebrospinal fluid (CSF) infusion test and PRx was calculated from CSF pressure and arterial pressure recordings. A simple scale was used to mark the patients’ outcome 6 months after surgery (improvement, temporary improvement, and no improvement).

**Results:** PRx was negatively correlated with R_out_ (R=−0.18; p=0.044); patients with normal CSF circulation tended to have worse autoregulation. The correlation for patients who were surgically-managed (N=83) was: R=−0.28; p=0.03, and stronger in patients who improved after surgery (N=64; R=−0.36; p=0.03). In patients who did not improve, the correlation was not significantly different from zero (N= 19; R=0.17; p=0.15). There was a trend towards higher values for PRx in non-responders than in responders (PRx =0.16+/− 0.04 vs 0.09 +/−0.02 respectively; p=0.061), associated with higher MAP values (107.2+/−8.2 in non-responders vs 89.5+/−3.5 in responders; p=0.195). The product of MAP* (1+PRx), proposed as a measure of combined arterial hypertension and deranged autoregulation, showed a significant association with outcome (greater value in non-responders; p=0.013).

**Conclusion:** Autoregulation proves to associate with cerebrospinal fluid circulation, and appears strongest in shunt responders. Outcome following CSF diversion is possibly most favorable when CSF outflow resistance is increased and global cerebral autoregulation is intact, in combination with arterial normotension.

## Introduction

Normal pressure hydrocephalus (NPH) is a neurodegenerative disease characterized by enlarged ventricles, and Hakim’s triad: gait disturbance, urinary incontinence, and dementia. Although prevalence estimates vary, it is known as the only “treatable dementia”, and thus warrants ongoing research into its pathophysiology, early diagnosis, and treatment ^1,2^. The pathophysiology of NPH, although better understood nowadays, remains to be elucidated. It can be divided in three components: disturbed CSF circulation, poor pressure-volume compensation and, possibly, abnormal Cerebral Blood Flow (CBF). Whether the regulatory mechanisms of CBF are disturbed cannot be concluded based on previous studies, primarily because almost every study has focused on demonstrating abnormal CBF patterns, which differ from our previous findings and current hypothesis on the importance of preserved autoregulation in outcome after shunting. Momjan et al (2004) showed reduced CBF in NPH patients white matter with a worsening pattern as the proximity to the ventricles increased. To the best of our knowledge, a few previous studies [Matthew et al, (1975 and 1977), Schmidt et al (1996) and Tanaka et al 1997) have correlated normal CBF before shunting with a positive outcome. Those studies, however, only relied on measurements of CBF alone and there are a lot more that contradicted this finding or were inconclusive on whether higher or lower CBF was associated with a better post-surgical outcome^1–5^. Other studies have indicated that the ability of the brain to maintain a constant CBF in response to changes in cerebral perfusion pressure (CPP) -termed cerebral autoregulation, may be of importance in the pathophysiology of NPH^6,7,18^. In support of this, patient outcome after shunting has been linked with preserved autoregulation pre-shunting^16,22,^; however, the association was rather weak and could not be replicated in a further study^21^.

The role of the CSF dynamics, characterized by resistance to CSF outflow (R_out_ and other pressure-volume compensatory parameters, is still ambiguous in NPH. Rout, calculated from routine CSF infusion tests, has been used as the most reliable infusion test -derived parameter in shunt response prediction. The road to the use of Rout as a diagnostic and outcome prediction tool was lead from early on with studies from Borgensen et al (1981 &1992) who reported a >80% post-shunting success rate for patients with Rout ≥13 mmHg*min/ml. The milestone dutch NPH trial^1^ provided the necessary evidence for the use of the most common threshold for Rout, 18 mmHg*min/ml, reporting a 92% success rate. This trial pioneered in quantitavely reporting clinical investigations for the patients before and after shunting, however those highly positive results could not be replicated ever since. The current knowledge on Rout and the different proposed thresholds for diagnosis in NPH are summarized on **Table 1**.

We previously reported a reciprocal relationship between the transcranial Doppler-based autoregulation index Mx and R_out_ in non-shunted patients under investigation for NPH, such that those with lower R_out_, had worse cerebral autoregulation^11^. We hypothesized that in patients with low R_out_ (suggesting normal CSF circulation), disturbed cerebral autoregulation may be in part responsible for the clinical symptoms. Similarly, in a cohort of both non-shunted and shunted patients, we found a reciprocal relationship between an intracranial pressure (ICP)-based marker of cerebral autoregulation (the cerebral pressure reactivity index-PRx) and R_out_^10^. Although these previous investigations direct us towards a negative correlation between R_out_ and brain autoregulation in NPH patients, the external validity of this relationship is limited by the relatively small sample sizes, and the relevance of the relationship with respect to patient outcome is unclear.

The objectives for this study were first to confirm the relationship between cerebrovascular function - as measured by the pressure reactivity index (PRx), and CSF dynamics and circulation (characterized by R_out_), and secondarily to examine how autoregulation (PRx) and systemic vascular dysfunction may relate to patient outcome following surgical management.

## Materials and Methods

### Patients

Patients attended the Hydrocephalus Clinic, at Cambridge University Hospital (CUH) between the years 2003–2015. We selected patients of age >30 years with possible NPH. The inclusion criteria were: patients presenting with radiological evidence of ventricular dilation, baseline ICP <18 mmHg, and at least two of the main NPH clinical elements, including gait difficulties, cognitive problems/ memory loss, or urinary incontinence. In addition, patients were never shunted previously and were undergoing diagnostic investigation with a computerized lumbar CSF infusion test^8^, which is a method approved by the National Institute of Clinical Excellence (NICE, UK, 2008; IPG263) regularly used to facilitate the diagnosis and management of NPH in our hospital. Studies without continuous blood pressure measurement were excluded, as were studies performed under general anesthesia, because of confounding influences of CSF dynamics^19^. Using the above criteria, we selected a cohort of 131 patients used for retrospective data analysis. All patients signed individual consent forms for the study and data, after gathering clinical information, were anonymized and analyzed as a form of routine clinical audit.

### Data acquisition

Continuous arterial blood pressure (ABP) was measured using a Finapress finger cuff (Ohmeda, Englewood, CO),^11^, while continuous waveforms of ICP (as indicated by lumbar CSF pressure) were obtained by connecting the lumbar puncture needle to a fluid filled pressure transducer (Edward Lifesciences™) and pressure amplifier (Spiegelberg or Philips). In all patients, adequate CSF pressure and arterial pulse waveforms were assured. All data and digital signals were recorded at 30–100 Hz and then processed using ICM+^®^ version 8.3 (University of Cambridge Enterprise Ltd)^33,34^

### Signal processing

Artifacts (movements, coughing, dislocations of finger blood pressure cuffs, etc.) were manually removed in every patient’s raw data file prior to analysis. The pulse amplitude (AMP) of ICP was calculated as the fundamental amplitude in the frequency band of 40 to 140 beats per minute. The magnitude of slow waves of ICP was quantified as the square-root of the power of the signal in the frequency band of 0.3–4 cycles per minute^8,9,19^.

The pressure reactivity index (PRx) was calculated as the correlation coefficient between 30 consecutive 10-second averages of mean ICP and mean ABP (MAP). To eliminate, as accurately as possible, the interference of gradually increasing ICP on the slow waves of ICP implicated in the PRx calculation, ICP was first detrended using a moving average high-pass filter [**Figure 1**]. The cerebrospinal compensatory index (RAP) was calculated similarly as the correlation between 30 consecutive 10-second averages of mean ICP and AMP^8,13,14^.

**Figure 1.**
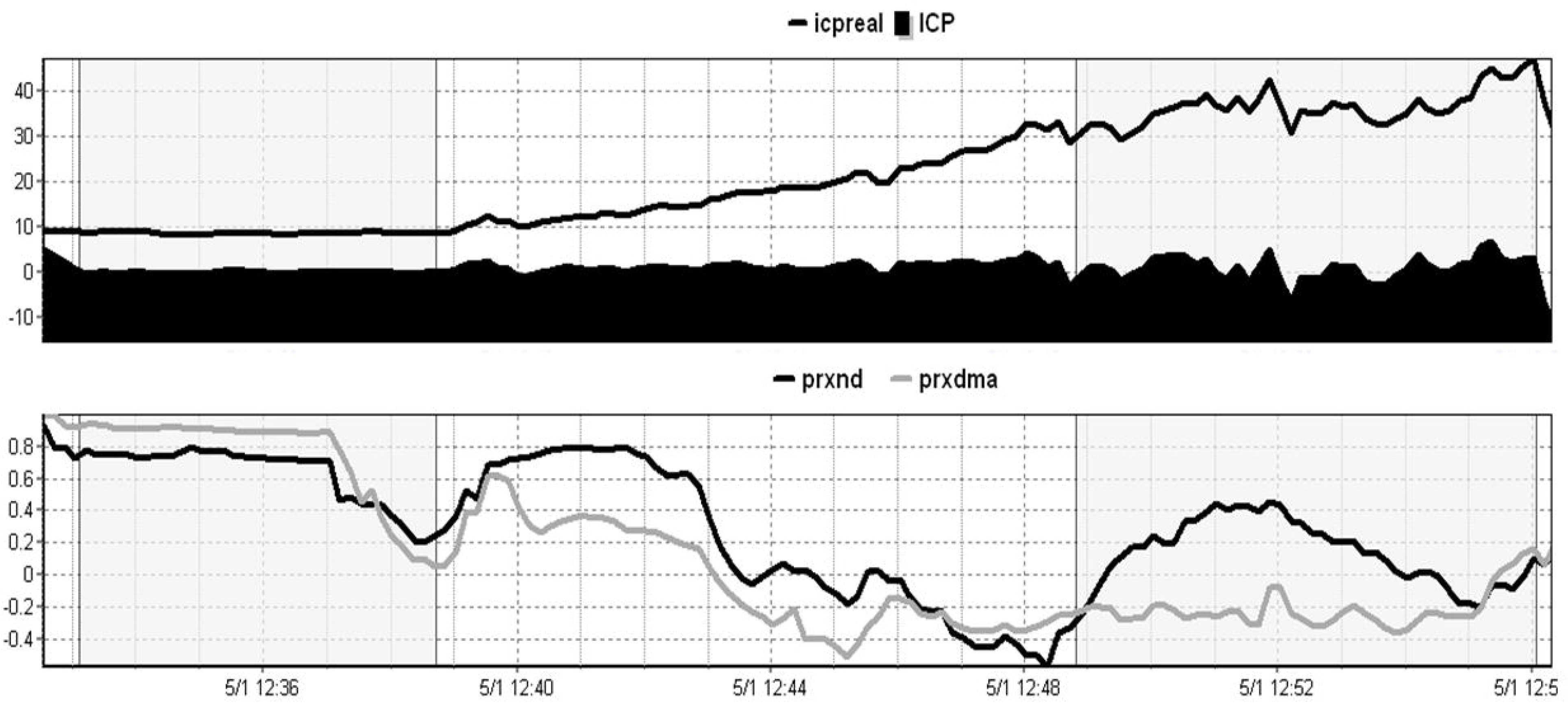
Difference in the trend between the original ICP and PRx (icpreal& prxnd respectively) and the detrended signal (ICP& PRx respectively); selected calculation periods of PRx at baseline and during icpreal: Original ICP singal, ICP: detrended ICP, prxnd: PRx non-detrended, prxdma: PRx detrended with moving average filter.

The aforementioned variables were averaged during the baseline and plateau phases of the infusion test as well as during the entire test (duration 25–40 minutes) [Figure 1].

The resistance to CSF outflow was calculated using a simple model integrated in our ICM+ software ^8,11^. The algorithm and the model used to calculate CSF resistance and elasticity has been reported in detail before and has been subject to modifications and improvements^8,11,13^. In some cases, strong vasogenic waves or elevation of the pressure above the safe limit of 40 mm Hg precludes precise measurement of the final pressure plateau. Computerized analysis produces results even in difficult cases in which the infusion is terminated prematurely. In order to avoid physiologically impossible and inaccurate measurement, only static R_out_ was used for further analysis ^34^. The slope of the ICP Amplitude – mean ICP line (AMP-P line) was determined by regression analysis of AMP and mean ICP using scatter plot charts provided as a function of ICM+^8,33^.

The product MAP* (1+PRx), produces a variable indicative of magnified vascular hypertension when autoregulation is impaired and attenuated vascular problems when autoregulation is preserved (PRx<0). As a concept similarly used before for ICP ^33^ we propose this product as a method that reflect the combined burden of hypertension and dysautoregulation in NPH patients.

### Patient follow-up

After having undergone lumbar infusion studies, patients were scheduled for follow-up in the hospital Hydrocephalus Clinic and a clinical decision on their further management was made. Patients and their families were carefully counseled and offered either a ventriculoperitoneal or ventriculoatrial shunt that included a programmable valve or an endoscopic IIIrd ventriculostomy (ETV). Outcome was determined from the hospital records. A simple scale, previously reported^25^, was used to classify patients who were shunted or underwent ETV, according to the clinical improvement of their symptoms after their operation. The scale classifies patient outcome as improved sustainably for 6 months (=1), improved for 3 months and then stopped improving or worsened (=2) and not improved after neither 3 nor 6 months (=3).

A subgroup analysis was performed to compare ETV patients with shunted patients based on both the studied parameters and the outcome after shunting. Chi-square test was used to compare the outcome in ETV vs shunted patients

### Statistical Analysis

After checking the normal distribution of data, Pearson’s correlation test was used to assess the correlation between the different parameters described above. Paired-samples t-tests were used to compare means and the difference between baseline and plateau measurements within the group. Independent samples t-tests were used to compare the means and the difference between in-cohort, separately defined groups of patients. Chi-square was used for qualitative data. Significance level was set at 0.05. Non-parametric tests (sign-ranked and Kruskal-Wallis) were used to compare results between the different outcome after shunting groups. All statistics were performed using the R software, version 3.3.1.

## Results

The mean age of the cohort 73 (+/−7) years and the male to female ratio was ~7:5 (77 males, 54 females). All of the patients except 8 had possible iNPH, without any clear etiology. Those 8 patients are the ones below the age of 55 and the NPH etiology was post-hemorrhagic, aqueductal stenosis, Chiari malformation (type II) or congenital hydrocephalus which had been treated in the past, had become shunt-independent and the patient had presented anew with symptoms resembling hydrocephalus.

CSF compensatory and autoregulatory parameters at baseline and during the infusion are presented in **Table 2** as mean values with standard errors of the mean, together with the level of significance (p-value) for change during the study. Notably, autoregulation, as expressed with PRx did not change during the test despite the fact that CPP decreased by 14.45 mmHg; p<0.01. Therefore, the mean PRx over the entire infusion study was used for subsequent correlation analysis.

### Global cerebral autoregulaton (PRx)

Overall, out of 131 patients, 30 showed disturbed autoregulation (PRx>0.25), 51 good autoregulation (PRx<0) and, in 50 patients, the autoregulation index fell within the inconclusive range (from 0 to 0.25). The numerical values for the other calculated parameters based on state of autoregulation (disturbed (PRx>0.25) and good autoregulation (PRx<0)) are given in **Table 3**. Increase in amplitude of ICP pulsation (dAMP) and compensatory reserve index RAP (dRAP) during the infusion studies were found to be significantly different between the autoregulating and non-autoregulating group.

### Interaction between PRx, R_out_ and other compensatory parameters

R_out_ was raised in 52 patients (>13 mm Hg*min/ml) and was within the ‘normal’ range (3.5 to 12.8 mm Hg*min/ml) in the remaining patients.

R_out_ was negatively correlated with PRx, rather weakly but significantly (R=−0.18; p=0.044) [**Figure 2**].

**Figure 2.**
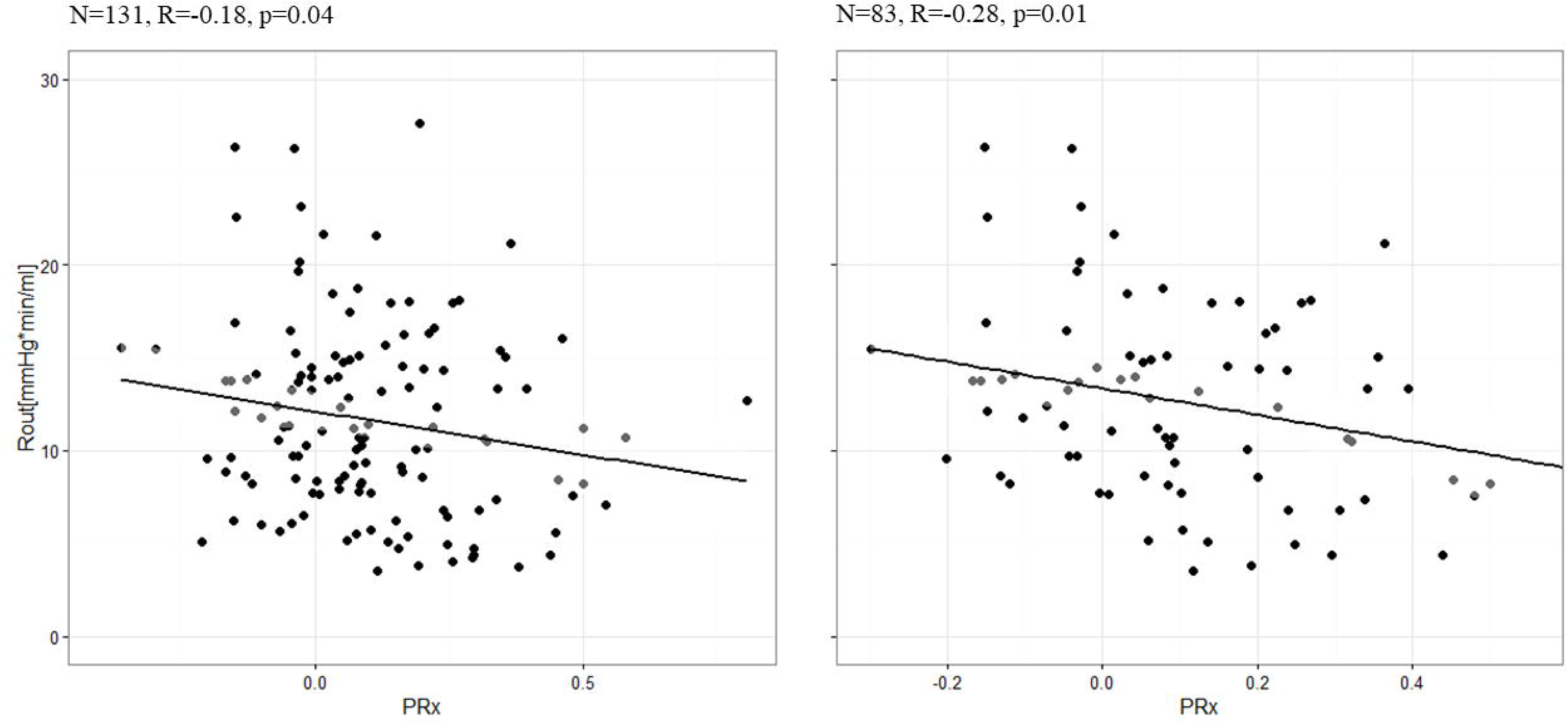
Left: Relationship between PRx and resistance to CSF outflow in our cohort of 131 non-shunted NPH patients undergoing lumbar infusion studies Right: Relationship between PRx and resistance to CSF outflow in patients who were clinically managed with ETV or shunted after the infusion studies.

Apart from autoregulation, R_out_ correlated with AMP (R=0.30; p<0.001). The correlation was even stronger between R_out_ and the increase in AMP during infusion (dAMP): R=0.46; p<0.001.

The correlation with PRx and other compensatory parameters (elasticity, ICP, baseline ICP, baseline AMP, slope of AMP/P line) was not significant. There was a significant correlation between the power of slow ICP waves and PRx (R=−0.19, p=0.025). PRx did not correlate with the age of patients (R =−0.09; p= 0.3119), while Rout was positively and relatively strongly correlated with patients’ age (R = 0.33; p = 0.00015).

### Interaction between surgery, PRx and Rout and impact on outcome

The clinical decision as to whether to offer CSF diversion surgery was based on clinical assessment, radiological image, and results of infusion studies plus other tests in the individual patient as part of overall assessment by the consultant neurosurgeon. The clinician was not aware of any of the other parameters studied, like PRx. Following the lumbar infusion study, 83 patients were clinically diagnosed with NPH and were either shunted (N=51), or underwent an ETV (N=32).

The negative correlation between PRx and R_out_ was significant when calculated for patients who were surgically-managed (N=83; R=−0.28; p=0.03) [**Figure 2**], whereas it was absent and not significantly different from 0 in the patients who were not offered CSF diversion (N=48, R= −0.05; p=0.7).

Following surgery, 64 patients improved at least temporarily, whereas 19 did not show any improvement. The relationship between R_out_ and PRx for the patients in the different outcome groups are demonstrated in Figure 3. A strong correlation was present in 64 patients who improved after surgery (N=64; R=−0.36; p=0.03). In 19 patients who did not improve with CSF diversion, the correlation was not significantly different from zero (N= 19; R=0.07; p=0.15).

**Figure 3.**
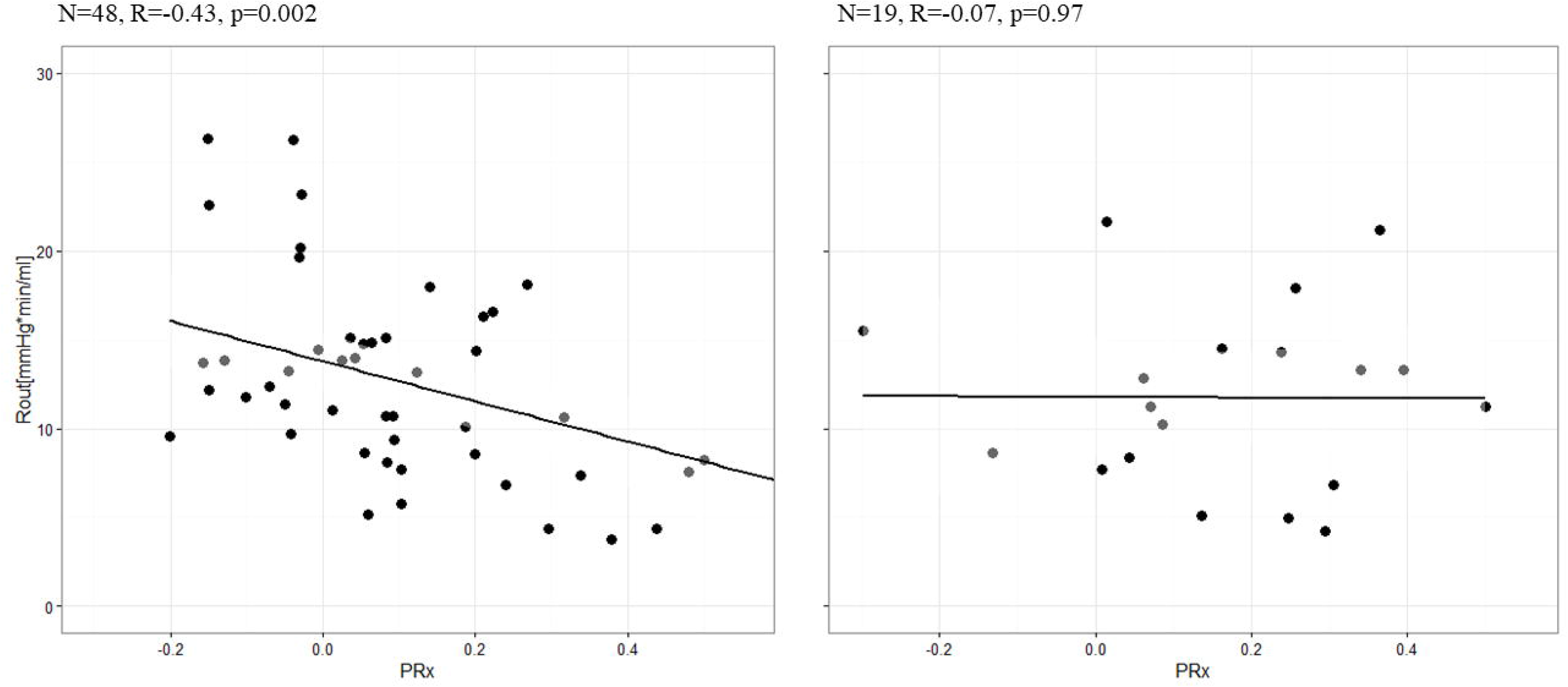
*Left: Relationship between PRx and resistance to CSF outflow in patients who improved sustainably after surgery (n=48, R=−0.43, p=0.002’). Right: Relationship between PRx and Rout in patients who did not improve after surgery (n=19, R=×0.007, p=0.97)*.

There was a trend, albeit not significant, towards higher values for PRx (more disturbed cerebral autoregulation) in non-responders than in responders to CSF diversion (PRx =0.16+/− 0.04 in nonresponders vs 0.09 +/−0.02 in responders; p=0.061) associated with higher (non-significantly) values for MAP (107.2+/−8.2 in non-responders vs 89.5+/−3.5 in responders; p=0.195). The product of MAP * (1+PRx), which is proposed as a measure of the combination of arterial hypertension and deranged autoregulation, was significantly associated with outcome (p=0.013). Comparisons of autoregulation and MAP parameters in the different outcome groups are further presented in **Table 4**, as mean values including standard errors of the mean. R_out_ was not different between responders and non-responders, however the group of studied patients was not powered to demonstrate such a difference.

### ETV vs shunted subgroup analysis

27 patients underwent ETV vs 56 who were shunted. The mean age for the ETV group was 55.74 +/− 3.19 vs 66.75 +/− 1.53 for the shunted group; p-value = 0.001625.

The mean AMP was different between the two groups (1.46 +/− 0.14 for ETV vs 2.41 +/− 0.16 for shunted; p=0.0005297). PRx was also different (0.17 +/−0.039 for ETV vs 0.07 +/− 0.02 for shunted; p-value = 0.012). Finally, Rout differed between ETV (12.0 +/− 1.34 mm Hg*min/ml and shunted patients (16.3 +/− 1.21 mmHg*min/ml; p-value = 0.003). When PRx was correlated with Rout in the ETV and the shunted groups separately, they did not correlate significantly in either groups (R= −0.27 p-value = 0.176 vs R=−0.2555274 p-value = 0.05733).

As to the outcome of ETV versus shunted patients, 21/27 patients after ETV had a positive outcome [≤2,] vs 43/56 shunted patients. 6/19 had a negative outcome after ETV vs 13/57 shunted individuals. Shunted patients did not show a significantly more positive outcome than ETV patients (xhi-square = 0.0102; p-value= 0.91.

## DISCUSSION

The relationship between the resistance to CSF outflow and pressure-reactivity index, although not very strong, was confirmed to be significant. Furthermore, an index of vascular function -as indicated by the combination of cerebral autoregulation and mean arterial pressure, was greater in patients with a lower chance of improving following shunt or ETV.

It is a very difficult endeavor to attempt utilizing two very conflicting aspects of NPH, Rout and autoregulation as a possible strong tool in NPH diagnosis and outcome prediction. It is unfortunate that after many years of very optimistic and polarly opposite, pessimistic views on both aspects there is uncertainty over those fundamental entities; the way forward from imaging and clinical scores appears inhibited. Etiology and pathophysiology of NPH need to be clarified as much as outcome prediction is in need of accuracy. Using ICP and autoregulation measurement techniques remains of the utmost importance in NPH until the achievement of optimal clinical management and outcomes; this includes many more studies to clarify and finally sort the ongoing discrepancies over definition, ambiguous diagnosis and usefulness of CSF dynamics parameters and role of the cerebrovascular bed^6,7,12^.

Tools such as infusion studies and ICM+ software have been the same since 2003^33,34^. We also have homogeneous criteria for admitting patients for hydrocephalus investigations in the CUH Hydrocephalus Clinic, introduced before 2003 and remaining unchanged. However, we cannot account in detail for the complete NPH assessment and referral process of our plethora of different neurosurgical consultants and this deviates from the initial purpose of our retrospective study.

High R_out_, despite criticism^1–5,10,25^, seems to be a major component of NPH, highlighting the importance of a disturbed CSF circulation ^1,5,15^. However, when R_out_ is used as a sole predictor of shunt response in large cohorts of patients, it fails to provide an accurate indication for surgery, as negative predictive power of R_out_ remains too low^1,36^. There is not enough scientific evidence to support the use of only R_out_ in the final clinical decision whether to opt for surgical intervention. A closer examination of PRx and the state of the individual’s autoregulation may therefore help fill this important gap in the decision-making for the management of NPH.

The criteria used by the consultants in our clinic were primarily based on careful counselling with the patient and family to explain balance of probable benefit and risk. First of all, the patients had to have radiological evidence of hydrocephalus and classical gait disturbance for any further consideration. The diagnosis and proposed CSF diversion were given definitely if the measure Rout from the infusion test was ≥ 13 mmHg*min/ml, If Rout was ≤ 13 mmHg*min/ml but there were minimal white matter changes on the MRI scan, the diagnosis and proposed surgical management were communicated as probable. However, after 2013, Extended lumbar drainage was implemented in our practice for those with an Rout ≤ 13 and, if there was a positive response to the drainage, a shunt was offered. A positive response was marked as an improvement in gait scores before and after the drainage, which seems the most prominent parameter improved during this time frame. Similarly, before 2013, the benefit for the patient was communicated as equivocal if Rout was ≤ 13 and the deep white matter lesions were very marked and the outcome prediction relied on the response to the CSF tap test performed after the infusion test (withdrawal of 30–50 mls). After 2013 those patients also underwent extended lumbar drainage, as described above^12,16,17^.

Our results agree with the previous reported findings suggesting that autoregulation is better retained in patients with abnormally increased R_out_^10,11^. This perhaps paradoxical relationship between autoregulation and R_out_, now replicated with a greater power, was hypothesized to be due to the presence of cerebrovascular disease in those patients presenting with a normal or low R_out_ ^11,16,18,21^. We did not possess further data to support this hypothesis at that time. However, we have now observed that when 83 of the patients studied were clinically diagnosed with hydrocephalus and underwent surgery, the correlation between disturbed autoregulation and lower R_out_ tended to be stronger. Even more interestingly, in the patients in whom the clinical decision was that they would most probably not benefit from surgery, no correlation between R_out_ and PRx was present. This finding could possibly suggest a distinction between patients who might benefit from a CSF diversion and whose clinical symptoms could be caused by disturbed CSF circulation, with or without cerebrovascular disease, and at least not by vascular disease alone. Autoregulation appears to have the potential to serve as a supplementary tool for clinical decision-making. The presence of vascular disease, both as differential and as comorbidity in hydrocephalus, constitutes one of the biggest hurdles in clinical practice and management^17,22,23,28^. There is still no alternative clinical test and consensus to aid the clinical decision whether there is concomitant disturbance of the CSF circulation and cerebrovascular disease, or whether the CSF circulation is normal and the problem lies purely in the vasculature^10,22,24^. PRx did not show any correlation with age, which is surprising, as significant correlation may be seen in anesthetized and ventilated patients after traumatic brain injury^9,20^. An additional similar correlation was shown between age and dysautoregulation in patients undergoing non-neurosurgical elective surgery. It appears that the age-autoregulation relationship (higher age associated with dysautoregulation) may be magnified by general anesthesia. On the other hand, R_out_ is known to positively correlate with age^8,15^. A strong relationship was successfully replicated in our cohort of NPH patients; a possible investigation of different Rout thresholds for different ages remains to be performed separately, for better future directions. Sex does not appear to have any particular influence on R_out_.

The CSF infusion test seems to provide ideal conditions for measurement of cerebral autoregulation using different methods. We demonstrated a significant decrease in CPP during infusion.^8,13^. Another autoregulation-associated parameter, the magnitude of slow waves, correlated weakly but significantly with PRx. Slow waves, in context of measurement of PRx or transcranial-Doppler investigation, are carriers of the information about autoregulation mechanism. This correlation has not been reported before and it is worthwhile to consider its role in NPH patients, since both the frequency and the magnitude of slow waves are involved in possibly distinguishing NPH patients without vascular problems and predicting their benefit from a shunt insertion^26–28^.

Furthermore, the correlation between R_out_ and PRx was found to be stronger in the patients that improved after shunting, and tended to be absent in the patients with no improvement. Outcome after shunting in NPH is very much multifactorial, and connecting it with one measured parameter is probably naïve^28–31^. This can be demonstrated by the finding that neither PRx nor R_out_ differed between the outcome groups but the correlation between PRx and R_out_ was found to differ significantly. PRx is a global autoregulation index, which means that it reflects the state of autoregulation in whole of the brain. It is a continuous index with low values meaning better autoregulation and high values denoting impaired autoregulation. Autoregulation in hydrocephalus is still an uncertain territory and this is the first time PRx was used in a large cohort of patients. The limits good autoregulation (<0), grey zone (0–0.25) and impaired autoregulation (>0.25), were determined in TBI patients and their applicability in hydrocephalus in unknown and needs to be studied. Over one third of the patients (62/131), had a PRx in the grey area: (PR =0−0.25) and we need more data in the future to determine the interpretation and the significance of this relationship with the Rout and with outcome. Our results create a new framework of consideration for patient outcome, moving further from the unsuccessful attempts to associate a single parameter strongly with shunt response; the addition of vascular disease markers and autoregulation indexes would be worth further investigating for outcome consideration in NPH.

It was not within the scope of this article to compare ETV versus shunting as a treatment option for hydrocephalus, nor did we possess enough patients to conclude this. The question of when ETV or shunt should be selected for different hydrocephalus patient remains to be tackled. However, there were some differences in PRx and Rout between ETV and shunted patients, that we cannot interprete at the moment but should be investigated separately at a larger cohort. The influence of those differences in our overall results does not appear significant by our current number of patients, at least as far as the correlation of PRx and Rout and outcome are concerned. Finally, as the guidelines on ETV and shunting in NPH have been ambiguous and the reasons for performing an ETV on our patients can vary, it is not easy to investigate this matter further as we are not able to report more information on the patients’ clinical course, from the first consultation to the final decision for surgery^11,35^.

Finally, it is important to point out that we presented a correlation between a lack of response to shunting and vascular (greater arterial pressure) as well as autoregulatory dysfunction combined. There is an abundance of data suggesting an interaction between hemodynamics and CSF circulation ^6,7,28–31,37^. Strengthening the significant but not quite ideal predictive value of R_out_ could prove valuable in clinical practice and spare patients from undergoing difficult, repetitive diagnostic procedures to determine their further management.

### Limitations

ABP was measured non-invasively using a Finapres finger cuff device, and while this has been shown to be accurate compared to invasive methods [17] its accuracy in this particular patient group is unclear. Furthermore, infusion studies have a limited duration and do not always allow optimal calculation of the pursued autoregulation indices which typically require long recordings. Longer recordings serve as more ideal examples but are not always possible and are dependent on patient cooperation.

We had not obtained any measurements of cerebral blood flow during the infusion studies. Also, PRx is an indirect index of cerebral autoregulation. As mentioned above, it appears that a significant proportion of the hydrocephalic patients studied have a “grey zone” PRx. More studies are needed to determine the limits of PRx for hydrocephalic patients and furthermore compare different methods of autoregulation assessment. Also, the MRI scans of the patients available could provide us with some rough evidence of possible vascular disease, however they were not available for some of the patients in our cohort as MRIs for possible NPH have not always been performed routinely in our hospital. A possible avenue for future MR-based correlation of vascular disease would be to assess for the burden of periventricular and deep white matter hyperintensitites to explore this relationship.

The patients referred for infusion studies come from referrals from a multitude of neurosurgical consultants. Due to the retrospective nature of our study and to its design, it was impossible and not in our goals to possess all the clinical information regarding the detailed spectrum of the patients’ disease, including the exact gait and memory scores. Based on evaluation from our experts, all of these patients were possible or probable NPH candidates undergoing further tests to ascertain the diagnosis and predict the outcome. Ventriculomegaly, gait impairment, dementia and urinary incontinence were diagnosed by specialists using the MMSE for cognitive assessment and gait scores from Tinetti Balance score and 10’ walking test but we are unable to report those exact scores.

Finally, the sample size we had available for correlating with different outcomes is relatively small, especially for the patients with poor outcome. New studies with a stronger sample are required in order to validate our results related to patients’ outcome.

## CONCLUSION

Our current study provides a basis to begin considering autoregulation in hydrocephalic patients. Further prospective trials should be conducted to elucidate the role of autoregulation to optimize management of NPH.

## Acknowledgement

JD is supported by a Woolf Fisher scholarship (NZ); JDP: NIHR Brain Injury HTC and NIHR Senior Investigator Award (2009–2014).

We thank Miss Leanne Calviello for language editing and proofreading.

